# Cholecystokinin-expressing GABA neurons elicit long-term potentiation in the cortical inhibitory synapses and attenuate sound-shock associative memory

**DOI:** 10.1101/2024.10.08.617188

**Authors:** Ge Zhang, Kwok Kin Pang, Xi Chen, Fengwen Huang, Jufang He

## Abstract

Neuronal interactions between inhibitory and excitatory neurons play a pivotal role in regulating the balance of excitation and inhibition in the central nervous system (CNS). Consequently, the efficacy of inhibitory/excitatory synapses profoundly affects neural network processing and overall neuronal functions. Here, we describe a novel form of long-term potentiation (LTP) induced at cortical inhibitory synapses and its behavioral consequences. We show that high-frequency laser stimulation (HFLS) of GABAergic neurons elicit inhibitory LTP (i-LTP) in pyramidal neurons of the auditory cortex (AC). The selective activation of cholecystokinin-expressing GABA (GABA^CCK^) neurons is essential for the formation of HFLS-induced i-LTP, rather than the classical parvalbumin (PV) neurons and somatostatin (SST) neurons. Intriguingly, i-LTP can be evoked in the AC by adding the exogenous neuropeptide CCK when PV neurons and SST neurons are selectively activated in PV-Cre and SST-Cre mice, respectively. Additionally, we discovered that low-frequency laser stimulation (LFLS) of PV neurons paired with HFLS of GABA^CCK^ neurons potentiates the inhibitory effect of PV interneurons on pyramidal neurons, thereby generating heterosynaptic i-LTP in the AC. Notably, light activation of GABA^CCK^ neurons in CCK-Cre mice significantly attenuates sound-shock associative memory, while stimulation of PV neurons does not affect this memory in PV-Cre mice. In conclusion, these results demonstrate a critical mechanism regulating the excitation-inhibition balance and modulating learning and memory in cortical circuits. This mechanism might serve as a potential target for the treatment of neurological disorders, including epilepsy and Alzheimer’s disease.

## Introduction

Memory formation and loss are primarily mediated by the modulation of synaptic plasticity in the central nervous system (CNS), which is considered a critical factor influencing the survival and lifespan of animals^1,2^. Long-term potentiation (LTP) and long-term depression (LTD) are the two major cellular mechanisms for modulating learning and memory in mammals^3,4^. Therefore, this research field has received tremendous attention from the neuroscientists over the past several decades^5,6,7^. Additionally, it has been well-documented that N-methyl-D-aspartate receptor (NMDAR) and α-amino-3-hydroxy-5-methyl-4-isoxazolepropionic acid receptor (AMPAR) play pivotal roles in modulating LTP and LTD, subsequently affecting the excitatory transmission between pre- and postsynaptic-neurons^8,9^. Notably, current optogenetic techniques combined with transgenic animal models are well developed for precise activation or inhibition of specific types of neurons^10,11,12^, facilitating researchers in uncovering more details about the neuronal processes underlying neuroplasticity and its influence on animal behavior.

Neuromodulators are broadly distributed and expressed in the central and peripheral nervous system, including serotonin (5-HT)^13^, dopamine (DP)^14^, and noradrenaline (NA)^15^, acetylcholine (ACh)^16^ and cholecystokinin (CCK)^17^. Importantly, these neuronal chemicals play critical role in shaping brain development and function in multiple aspects, including neuroplasticity and its behavioral effects^13, 14, 15, 16, 17^. CCK, as the most abundant neuropeptides in the CNS, has been implicated in regulating various physiological and neurobiological states^18,19^. Significant advancements have been made in understanding the role of CCK in the brain functions^19,20,21,22,23^.

For instance, previous studies have demonstrated that entorhinal cortex derived CCK (EC^CCK^) facilitates hetero-synaptic LTP formation in the auditory cortex (AC) ^19^, bilateral amygdala (BLA)^20^ and dorsal hippocampus (DHP) ^17^, subsequently modulating the sound-sound associative memory, trace fear memory and spatial memory, respectively. Intriguingly, recent study have further demonstrated that selective activation of GABA^CCK^ synapses significantly and permanently potentiates inhibitory postsynaptic currents (IPSC) in pyramidal neurons of the AC. This form of neuroplasticity, known as inhibitory LTP (i-LTP), is mediated by the novel CCK-receptor (GPCR-173) rather than CCK-1R or CCK-2R^21^. However, it remains unclear whether other interneurons are involved in regulating i-LTP formation in the AC and how changes in synaptic plasticity in the AC affect subsequent behavioral performance. Further investigations are needed to elucidate how endogenous neuropeptide CCK interacts with these interneurons regarding neurobiological functions in the AC.

To address these questions, we combined multiple experimental approaches in the present study, including optogenetic manipulation, whole-cell recording, immunohistochemistry, transgenic cross-breeding, and behavioral paradigms. These methods were used to systematically elucidate the differences in neuronal processes between GABA^CCK^ neurons and other interneurons in shaping neuroplasticity in the AC and its behavioral consequences. Ultimately, our study unveiled a novel mechanism by which GABA^CCK^ neurons modulate neurobiological functions in cortical circuitry at both electrophysiological and behavioral levels.

## Results

### HFLS of GABA neuron induced i-LTP on pyramidal neuron in the AC

The neuronal and cellular mechanisms of long-term potentiation (LTP) mediated by the excitatory neuromodulator (e-LTP) have been extensively investigated^5,6,7^. However, the role of inhibitory mechanisms in modulating LTP via optogenetics remains poorly understood (**Figure A**). Thus, we utilized GABA-Cre mice and Cre-dependent AAV containing the light-sensitive channelrhodopsin (Chronos) to examine the neuronal role of neurotransmitter GABA in inhibitory synapses. To specifically target GABA neurons, we injected the AAV9-mDlx-DIO-Chronos-mCherry into the auditory cortex (AC) of GABA-Cre mice **(Figure 1B-1C)**. Four weeks after AAV expression. most mCherry^+^ neurons exhibited colocalization with the GABA marker (**Figure 1D**).

**Figure 1.**
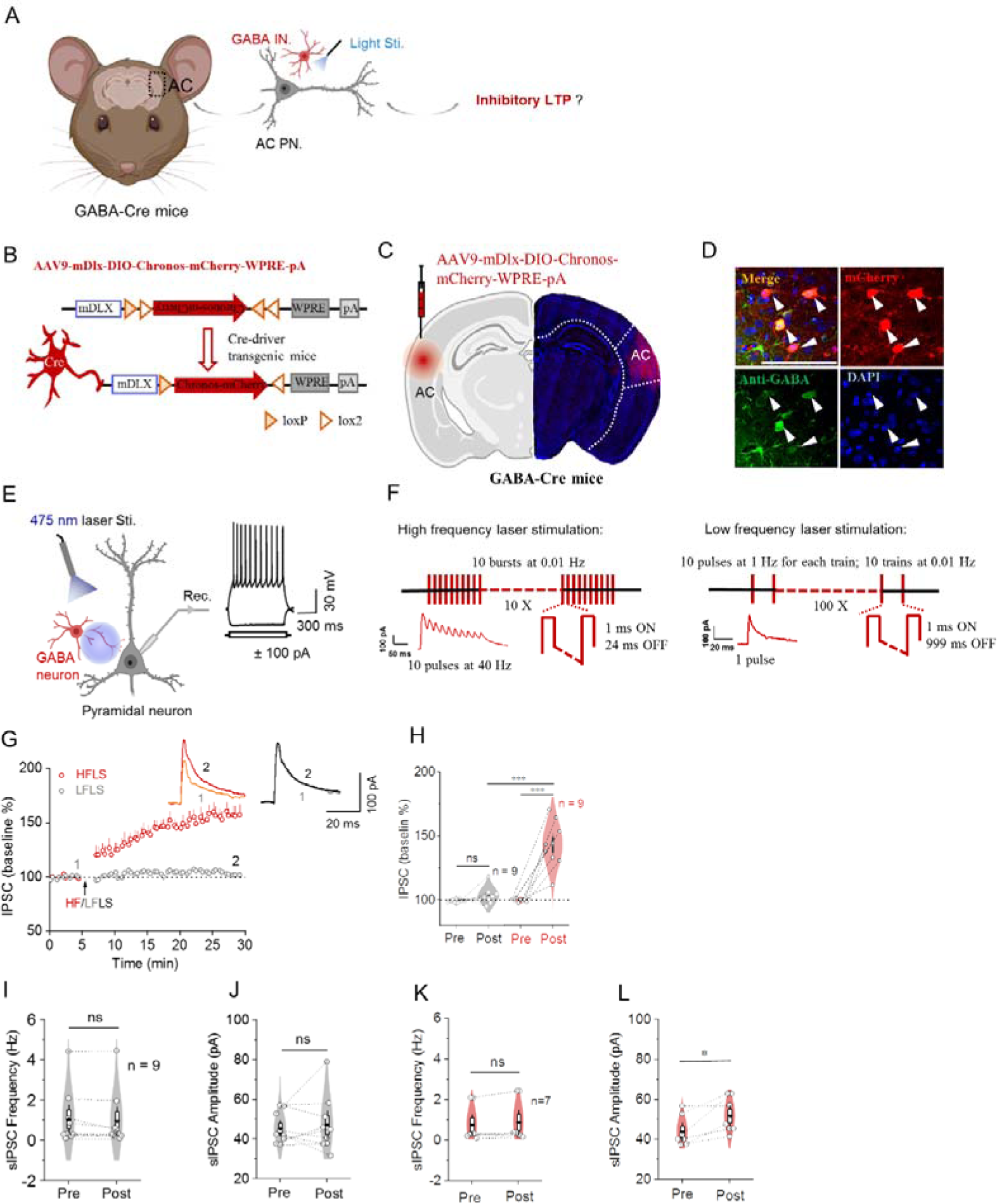
HFLS of GABA neuron induced i-LTP on pyramidal neuron in the AC. **(A)** Schematic demonstration of how GABA neurons exhibit its neuronal effects on pyramidal neurons in auditory cortex (AC PN.). **(B)** Schematic illustration of Cre-dependent AAV vector (DIO-Chronos-mCherry) injection into the AC of GABA-Cre mice to specifically infect GABA neurons. **(C)** Schematic diagram of virus injection into the AC of GABA-Cre mice (AAV9-mDlx-DIO-Chronos-mCherry-WPRE-pA, 6.15E+12 vg/mL, 200 nL for two different sites). **(D)** Confocal image of virus expression in the AC. White arrows indicate the colocalization of antibody-GABA with mCheryy expressed GABA positive neurons. Scale bar: 50 µm. **(E)** Schematic depiction of the IPSC recording of the GABA neuron in the AC (left); A representative trace of the regular spiking of a pyramidal neuron (right). **(F)** Protocols (HFLS and LFLS) for stimulating the Chronos expressed GABA neuron during the IPSC recording. **(G)** Normalized amplitude of IPSC before and after 473 nm wavelength LFLS (grey) and HFLS (red). **(H)** Statistical comparison of amplitudes of IPSC before and after 473 nm wavelength LFLS (grey) and HFLS (red). **(I)** Quantitative analyses of sIPSC frequency before and after the 473 nm wavelength LFLS. **(J)** Quantitative analyses of sIPSC amplitude before and after the 473 nm wavelength LFLS. **(K)** Quantitative analyses of sIPSC frequency before and after the 473 nm wavelength HFLS. **(L)** Quantitative analyses of sIPSC amplitude before and after the 473 nm wavelength HFLS. ^∗^p < 0.05, ^∗∗^p < 0.01, ^∗∗∗^p < 0.001; ns, not significant. Data are reported as mean ± SEM.

Subsequently, we used a 473 nm wavelength light to active the Chronos-expressing GABA neurons and recorded the inhibitory postsynaptic currents (IPSCs) in pyramidal neurons in the AC area while holding the membrane potential at −50 mV (**Figure 1E**). After establishing a stable baseline, we applied the high-frequency light stimulation (HFLS) to the GABA synapses (**Figure 1F**, left panel), which is presumed to trigger the release of neuromodulators from the vesicles in the pre-synaptic terminals^17^. Intriguingly, robust and permanent inhibitory LTP (i-LTP) was evoked in pyramidal neurons (**Figure 1G-1H**, two way ANOVA, Bonferrori adjustment, F_1,16_ = 52.65, p < 0.001; HFLS: Pre 99.66 ± 0.41 % vs Post 143.02 ± 5.99 %, p < 0.001; LFLS: Pre 99.24 ± 0.47 % vs Post 103.10 ± 2.27 %, p = 0.41; Post_HFLS_ vs Post_LFLS_, p < 0.001). In comparison, low-frequency laser stimulation (LFLS; **Figure 1F**, right panel) of GABA neurons did not result in significant changes in IPSCs. These results indicate that HFLS markedly reinforces the synaptic plasticity between inhibitory synapses and their corresponding pyramidal neurons.

Furthermore, the spontaneous IPSCs (sIPSCs) were also obtained from pyramidal neurons by holding the cells at 0 mV (S**Figure 1A-1B**). No significant differences were found in either the amplitude or frequency of sIPSCs before and after LFLS of GABA neurons (**Figure 1I**, paired t-test, t_8_ = −1.54, p = 0.16; Pre 1.06 ± 0.47 Hz vs Post 0.96 ± 0.47 Hz; **Figure 1J**, paired t-test, t_8_ = −0.51, p = 0.62; Pre 45.51 ± 2.64 pA vs Post 47.35 ± 4.76 pA). By contrast, HFLS notably potentiated the amplitude of sIPSCs (**Figure 1K**, paired t-test, t_6_ = −3.26, p = 0.02; Pre 43.78 ± 2.59 pA vs Post 51.80 ± 2.99 pA) and thereby markedly enhances the AP firing probability^24,25^. Concurrently, no significant changes in frequency of sIPSC by the same manipulation (**Figure 1L**, paired t-test, t_6_ = −2.10, p = 0.08; Pre 0.76 ± 0.30 Hz vs Post 0.89 ± 0.35 Hz).

### CCK-GABA neurons are crucial for HFLS-induced i-LTP

GABAergic interneurons constitute approximately 25-30% of the cortical neurons and play a crucial role in modulating cortical efferent and plasticity^26^. These interneurons can be categorized into different types, such as PV interneurons, somatostatin (SST) interneurons, vasoactive intestinal peptide (VIP) interneurons, CCK interneurons, and others, based on various neurological characteristics including electrophysiological properties, molecular expression profiles, and developmental origin^27^. Our previous study demonstrated that HFLS of glutamatergic entorhinal cortex-derived CCK afferents (EC^CCK^-AC projections) enables the formation of hetero-synaptic LTP on pyramidal neurons in AC area^19^. However, the neuronal effect of local GABA^CCK^ neurons on pyramidal neurons is still not well-understood (**Figure 2A**). Therefore, we further utilized various transgenic mice to unveil the role of different interneurons in regulating the neuroplasticity. Here, we primarily focus on the influence of CCK, PV and SST interneurons on i-LTP formation in AC, since the neural functions of PV and SST are relative well documented in AC area ^28,29,30^.

**Figure 2.**
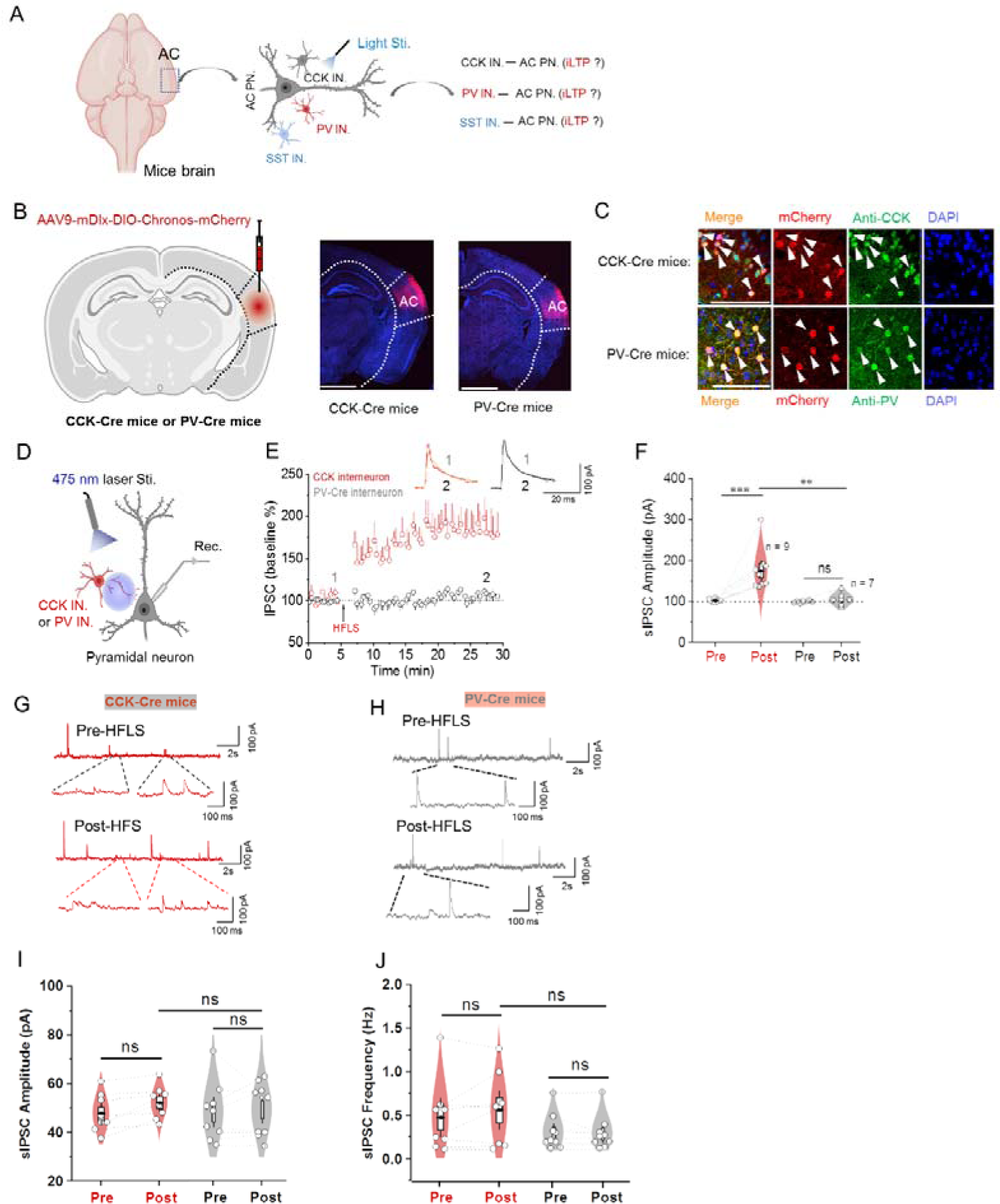
CCK-GABA neurons are crucial for HFLS-induced i-LTP. **(A)** Schematic demonstration of which type of interneuron contribute to the iLTP formation in pyramidal neurons of auditory cortex (AC PN.). **(B)** Schematic illustration of Cre-dependent AAV vector (DIO-Chronos-mCherry; 6.15E+12 vg/mL, 200 nL for two different sites) injection into the AC of CCK-Cre mice and PV-Cre mice to specifically infect GABAergic neurons, respectively. **(C)** Confocal image of virus expression in the AC of CCK-Cre mice (upper) and PV-Cre mice (bottom). White arrows indicate the colocalization of antibody-CCK/PV with mCheryy expressed CCK/PV positive neurons. Scale bar: 50 µm. **(D)** Schematic depiction of the IPSC recording of GABA^CCK^/PV interneuron in the AC. **(E)** Normalized amplitude of IPSC before and after 473 nm wavelength HFLS in CCK-Cre mice (red) and PV-Cre mice (grey), respectively. **(F)** Statistical comparison of amplitude of IPSC before and after 635 nm wavelength HFLS in CCK-Cre mice (red) and PV-Cre mice (grey), respectively. **(G)** Representative sIPSC traces of GABA^CCK^ neurons before (upper) and after (bottom) HFLS in the CCK-Cre mice. **(H)** Representative sIPSC traces of PV interneurons before (upper) and after (bottom) HFLS in the PV-Cre mice. **(I)** Quantitative analyses of sIPSC frequency before and after the 473 nm wavelength HFLS in CCK-Cre mice (red) and PV-Cre mice (grey), respectively. **(J)** Quantitative analyses of sIPSC amplitude before and after the 473 nm wavelength LFLS in CCK-Cre mice (red) and PV-Cre mice (grey), respectively ^∗^p < 0.05, ^∗∗^p < 0.01, ^∗∗∗^p < 0.001; ns, not significant. Data are reported as mean ± SEM.

Next, we injected AAV expressing DIO-mDLX-Chronos into the AC of CCK-Cre, PV-Cre mice and SST-Cre mice, respectively (**Figure 2B**; **SFigure 1A**). Four weeks after the AAV expression, most virus labeled inhibitory neurons were positive for their corresponding markers (**Figure 2C**; **SFigure 2B**). Additionally, IPSCs were recorded IPSC on pyramidal neurons by activating these interneurons expressing Chronos (**Figure 2D**; **SFigure 2C**). Interestingly, we observed a significant increase in IPSC amplitude in CCK-Cre mice following HFLS manipulation, while no significant IPSC potentiation was found in PV-Cre and SST-Cre mice under the same protocol (**Figure 2E-2F**, two way ANOVA, Bonferrori adjustment, F_1,14_ = 11.34, p = 0.005; HFLS: Pre 102.15 ± 1.21 % vs Post 175.70 ± 17.10 %, p < 0.001; LFLS: Pre 99.10 % ± 0.98 % vs Post 104.87 ± 5.12 %, p = 0.71; Post_CCK_ vs Post_PV_, p = 0.003; **SFigure 2D-2E**, paired t-test, t_9_ = 0.56, p = 0.59; Pre 100.34 ± 0.36 % vs Post 99.03 ± 2.52 %; **SFigure 2F**, one way ANOVA, Bonferrori adjustment; F_2,23_ = 16.30, p < 0.001; Post_CCK_ vs Post_SST_, p = 0.001). These findings suggest that GABA^CCK^ neurons are essential for eliciting i-LTP in the AC area.

Furthermore, spontaneous IPSC (sIPSC) events were also recorded from pyramidal neurons in the three groups of mice (**Figure 2G-2H; SFigure 2G**). However, HFLS of GABA^CCK^ neurons did not enhance the amplitude of sIPSC in CCK-Cre mice, as well as in PV-Cre mice and SST-Cre mice (**Figure 2I**; two way ANOVA, Bonferrori adjustment, F_1,15_ = 1.54, p = 0.23; CCK-Cre: Pre 47.92 ± 2.80 pA vs Post 52.42 ± 2.40 pA, p= 0.10; PV-Cre: Pre 48.51 ± 4.38 pA vs Post 49.53 ± 3.95 pA, p = 0.68; Post_CCK_ vs Post_PV_, p = 0.53; **SFigure 2H**, paired t-test, t_9_ = 1.97, p = 0.07; Pre 48.22 ± 3.27 pA to Post 44.50 ± 2.00 pA), suggesting that potentiated amplitude of sIPSC in GABA neurons is possibly mediated by the interactions of multiple neuromodulators rather than a single one. Concurrently, this manipulation also did not affect the frequency of sIPSC in any of the three groups (**Figure 2J**; two way ANOVA, Bonferrori adjustment, F_1,15_ = 1.54, p = 0.23; CCK-Cre: Pre 0.48 ± 0.15 Hz vs Post 0.56 ± 0.14 Hz, p = 0.14; PV-Cre: Pre 0.30 ± 0.07 Hz vs Post 0.29 ± 0.07 Hz, p = 0.87; Post_CCK_ vs Post_PV_, p = 0.09; **SFigure 2I**, paired t-test, t_9_ = 0.13, p = 0.9, Pre 0.35 ± 0.06 Hz to Post 0.34 ± 0.08 Hz). Overall, these results unveil a unique role of CCK in modulating the HFLS-induced iLTP in the AC area.

### Exogenous CCK facilitates the formation of i-LTP in the AC

Based on the results presented in **Figure 2E-2F**, it is evident that HFLS of GABA^CCK^ synapses notably potentiates inhibitory input on the pyramidal neurons rather than PV or SST interneurons. This enhancement is likely due to the release of endogenous CCK from the presynaptic terminals, leading to the formation of i-LTP under these conditions. To verify this hypothesis, exogenous CCK-8s was introduced into the ACSF solution during HFLS of the synapses in PV-Cre and SST-Cre mice (**Figure 3A**). Consistent with previous experimental procedures, following a stable baseline recording, the ACSF solution containing CCK (400 nM) was perfused, and the HFLS protocol was applied (**Figure 3B)**. Intriguingly, a progressive increase in inhibitory postsynaptic currents (IPSC) was observed in the pyramidal neurons of PV-Cre mice (**Figure 3C-3D**, two way ANOVA, Bonferrori adjustment; F_1,12_ = 3.14, p = 0.10; HFLF + CCK8s: Pre 100.26 ± 0.50 % vs Post 142.86 ± 4.13 %, p < 0.001). In contrast, LFLS paired with CCK also elicited comparable i-LTP (LFLF + CCK-8s: Pre 100.79 ± 0.72 % vs Post 130.45 ± 5.75 %, p < 0.001; Post_HFLF + CCK-8s_ vs Post _LFLF + CCK-8s_, p = 0.10). This finding suggests that the coordinated activation of inhibitory synapses, along with the neuronal peptide CCK, is crucial for the establishment of i-LTP.

**Figure 3.**
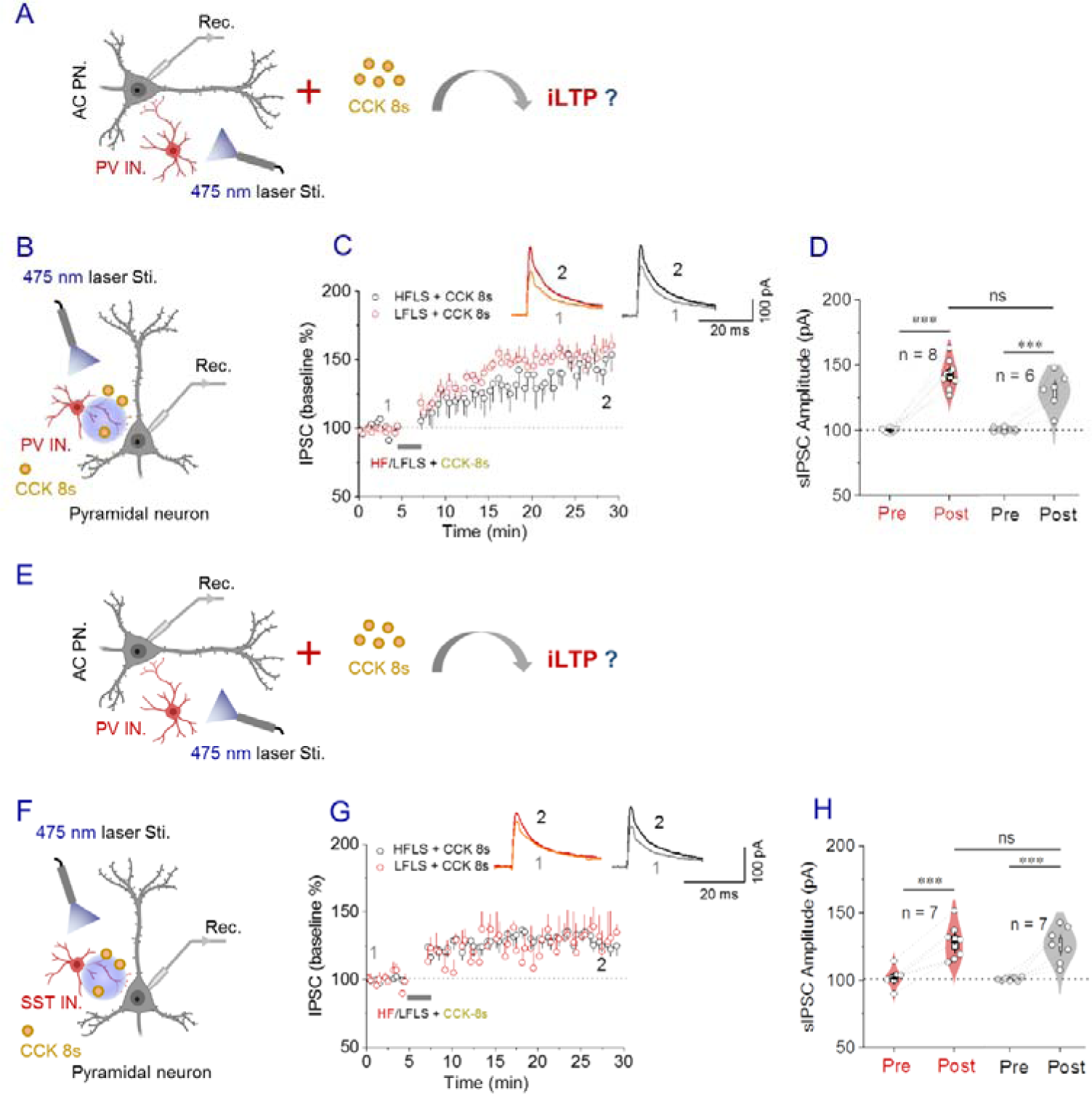
Exogenous CCK facilitates the formation of i-LTP in the AC. **(A)** Schematic demonstration of whether exogenous CCK interacts with PV interneuron to elicit iLTP in pyramidal neurons of auditory cortex (AC PN.). **(B)** Schematic depiction of the IPSC recording of PV interneuron in the AC of PV-Cre mice. **(C)** Normalized amplitude of IPSC before and after perfusion with CCK-8s in HFLS group (red) and LFLS group (grey), respectively. **(D)** Statistical comparison of the amplitude of IPSC before and after perfusion with CCK-8s in both groups. **(E)** Schematic demonstration of whether exogenous CCK interacts with SST interneuron to elicit iLTP in pyramidal neurons of auditory cortex (AC PN.). **(F)** Schematic depiction of the IPSC recording of the SST interneuron in the AC of the SST-Cre mcie. **(G)** Normalized amplitude of IPSC before and after perfusion with CCK-8s in HFLS group (red) and LFLS group (grey), respectively. **(H)** Statistical comparison of the amplitude of IPSC before and after perfusion with CCK-8s in both groups. ^∗^p < 0.05, ^∗∗^p < 0.01, ^∗∗∗^p < 0.001; ns, not significant. Data are reported as mean ± SEM.

Furthermore, the same protocol was employed to examine the neuronal effect of exogenous CCK-8s on pyramidal neurons in the AC of SST-Cre mice (**Figure 3E**). Similarly, a robust and sustained IPSC was elicited by exogenous CCK-8s (**Figure 3F-3H**, two-way ANOVA, Bonferrori adjustment; F_1,12_ = 0.02, p = 0.90; HFLF + CCK8s: Pre 101.86 ± 2.75 % vs Post 128.51 ± 5.13 %, p < 0.001). Additionally, equivalent potentiation of IPSC was found in the LFLS group (LFLF + CCK8s: Pre 100.69 ± 0.48 % vs Post 126.50 ± 5.00 %, p < 0.001; Post_HFLF + CCK8s_ vs Post _LFLF + CCK8s_, p = 0.78). Moreover, it was noted that CCK-mediated i-LTP in PV-Cre mice was slightly higher than in SST-Cre mice, although the reasons for this phenomenon remain unclear. One possibility is that post-modulation activities via CCK-8s are more pronounced in PV-Cre mice compared to SST-Cre mice. Based on these findings, we can conclude that neuronal CCK effectively regulates neuroplasticity mediated by both types of interneurons and facilitates the persistent influence of efferent neural responses on postsynaptic sites.

### Potentiation of inhibition effects of PV-interneuron on pyramidal neuron requires high-frequency activation of GABA^CCK^ neuron

Based on the aforementioned investigation into the effects of exogenous CCK on i-LTP formation, we aimed to determine whether HFLS of GABA^CCK^ neurons could enhance IPSC mediated by other interneurons in the AC area (**Figure 4A**). To achieve this, we utilized two different light-sensitive channelrhodopsins (Chronos and ChrimsonR) to selectively target GABA^CCK^ neurons and PV interneurons in genetically modified mice^18^.

**Figure 4.**
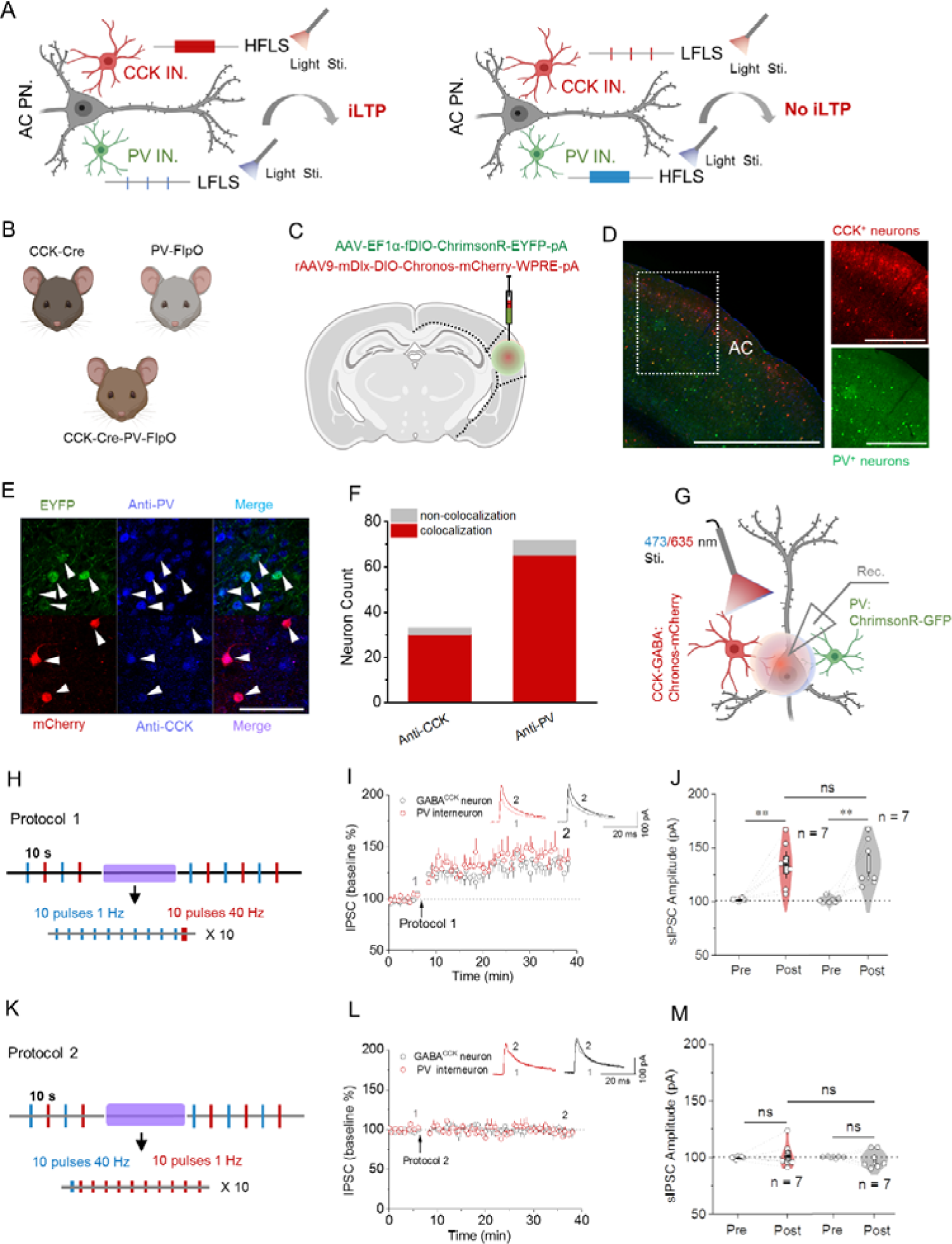
Potentiation of inhibition effects of PV-interneuron on pyramidal neuron requires high-frequency activation of GABA^CCK^ neuron. **(A)** Schematic demonstration of whether HFLS of CCK interneurons or PV interneurons facilitate the iLTP formation in AC PN. **(B)** Schematic illustration of the acquisition of CCK-Cre-PV-FlpO pups by crossing CCK-Cre mice with PV-Flop mice. **(C)** Mixture of AAV9-mDlx-DIO-Chronos-mCherry-WPRE-pA and AAV9-fDIO-ChrmosonR-EYFP-WPRE-pA was injected into the AC area of CCK-Cre-PV-FlpO mice (DIO-Chronos-mCherry/-fDIO-ChrimsonR-EYFP-pA; 6.15E+12 vg/mL, 200 nL for two different sites). **(D)** Fluorescent image showing AAV expression in AC area of CCK-Cre-PV-FlpO mice. **(E)** Confocal image of two kinds of AAV expression in pyramidal neurons in AC area. White arrows indicate the colocalization of antibody-CCK/antibody-PV with mCheryy expressed CCK and PV positive neurons, respectively. Scale bar: 50 µm. **(F)** Schematic depiction of two kinds of lights to separately evoke the IPSC in Chronos expressed GABA^CCK^ neuron and ChrimsonR infected PV neuron in the AC area of CCK-Cre-PV-FlpO mice. **(G)** Protocol consist of 473 nm HFLS and 635 nm LFLS for activating the Chronos expressed GABA^CCK^ neuron and ChrimsonR infected PV neuron, respectively. **(H)** Normalized amplitude of IPSC in GABA^CCK^ neuron and PV neuron before and manipulating the protocol 1, respectively. **(I)** Statistical comparison of the amplitude of IPSCs before and after manipulating the protocol 1 in both groups. **(J)** Protocol consist of 473 nm LFLS and 635 nm HFLS for activating the Chronos expressed GABA^CCK^ neuron and ChrimsonR infected PV neuron, respectively. **(K)** Normalized amplitude of IPSC in GABA^CCK^ neuron and PV neuron before and manipulating the protocol 2, respectively. **(L)** Statistical comparison of the amplitudes of IPSCs before and after manipulating the protocol 2 in both groups. ^∗^p < 0.05, ^∗∗^p < 0.01, ^∗∗∗^p < 0.001; ns, not significant. Data are reported as mean ± SEM.

Firstly, we crossed CCK-Cre mice with PV-FlpO mice to obtain CCK-Cre-PV-FlpO pups (**Figure 4B**). Heterozygous mice containing both Cre and FlpO elements were able to identify and separately infect CCK-positive and PV-positive neurons using Cre and FlpO-dependent AAV, respectively. Subsequently, we injected AAV9-mDlx-DIO-Chronos-mCherry-WPRE-pA and AAV9-fDIO-ChrmosonR-EYFP-WPRE-pA into the AC area of CCK-Cre-PV-FlpO mice to express Chronos-mCherry in GABAergic CCK neurons and express ChrimsonR-EYFP in PV interneurons, respectively (**Figure 4C**). Immunohistochemistry (IHC) analysis confirmed high colocalizations between the interneuron markers (anti-PV and anti-CCK) and their corresponding target neurons (PV and GABA^CCK^) in brain sections of CCK-Cre-PV-FlpO mice (**Figure 4D-F**). After a 4-week period of AAV expression, two different wavelengths of light (473 nm and 635 nm) were used to evoke Chronos-mediated IPSC in GABA^CCK^ neurons and ChrimsonR-mediated IPSC in PV neurons in the target region (**Figure 4G**).

For baseline recording, the membrane potential was held at −50 mV, and single-pulse light-evoked IPSC in GABA^CCK^ interneuron and PV interneuron were recorded using blue and red light interstimulus intervals of 10 s, Subsequently, HFLS pairs of 473 nm and 635 nm were applied to the target neurons. Intriguingly, this protocol significantly increased both GABA^CCK^-IPSC and PV-IPSC in AC pyramidal neuron (**Figure 4H-J**, two way ANOVA, Bonferrori adjustment, F_1,12_ = 0.01, p = 0.92; GBBA^CCK^: Pre 101.65 ± 0.49 % vs Post 134.51 ± 8.10 %, p = 0.002; PV: Pre 101.18 ± 1.11 % vs Post 135.32 ± 8.23 %, p = 0.001; Post_GBBA-CCK_ vs Post_PV_, p = 0.95), implying that HFLS to CCK^GABA^ neurons enhances the inhibitory effects of PV interneurons on pyramidal neurons, leading to the potentiation of neuroplasticity in a heterosynaptic manner (heterosynaptic-LTP). Additionally, another protocol (473 nm LFLS pairs with 635 nm HFLS) unsuccessfully elicit the heterosynaptic-LTP in AC area (**Figure 4K-M**, two way ANOVA, Bonferrori adjustment, F_1,12_ = 0.53, p = 0.48; GABA^CCK^: Pre 99.88 ± 0.38 % vs Post 101.79 ± 4.18 %, p = 0.61; PV: Pre 100.23 ± 0.31 % vs Post 98.38 ± 3.00 %, p = 0.62; Post_GBBA-CCK_ vs Post_PV_, p = 0.53). this finding indicates that heterosynaptic-LTP can be induced by HFLS of GABA^CCK^ neurons rather than PV interneurons in the AC region. Collectively, these findings reveal the essential role of neuronal GABA^CCK^ neurons in modulating neuroplasticity in the mammalian brain.

### HLFS of GABA^CCK^ neurons attenuate the sound-shock associative memory

Since neuroplasticity plays a critical role in modulating learning and memory in mammals, we conducted an investigation to determine whether activation of GABA^CCK^ neurons affects behavioral outcomes (**Figure 5A**). To test this assumption, we designed a behavioral paradigm to assess the animal’s response to sound-shock associative memory when GABA^CCK^ neurons were stimulated with light (**Figure 5B**). In brief, on the first day, mice were placed in a square apparatus equipped with foot-shock and audio devices. They were allowed one minute of free movement, followed by a tone cue and electrical stimulus. After a one-hour period for memory consolidation, light stimulation was administered to the trained mice. The following day, the mice were placed in a novel cylindrical apparatus and received a tone cue to assess freezing response (**Figure 5C**).

**Figure 5.**
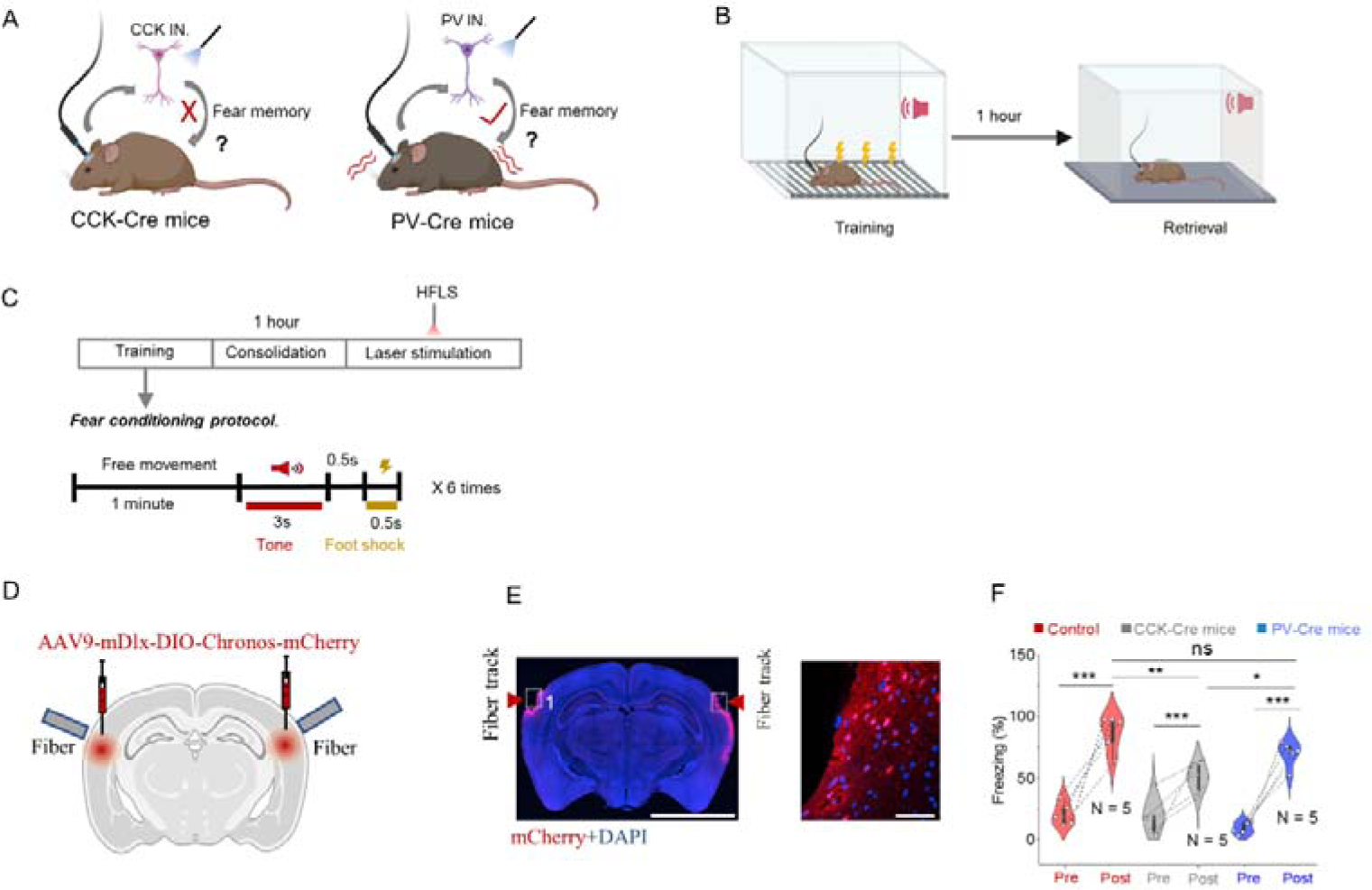
Potentiation of inhibition effects of PV-interneuron on pyramidal neuron requires high-frequency activation of GABA^CCK^ neuron. **(A)** Schematic demonstration of whether HFLS of CCK interneurons or PV interneurons affects animals’ fear memory. **(B)** Schematic depiction of behavioral diagram of sound-shock association memory. **(C)** The drawing of the protocol of the behavioral diagram. **(D)** Schematic illustration of AAV injection and fiber implantation in AC of experimental mice. **(E)** Fluorescent image showing the AAV expression and fiber track in AC area. **(F)** Statistical analysis of the freezing proportion in three different groups of mice. ^∗^p < 0.05, ^∗∗^p < 0.01, ^∗∗∗^p < 0.001; ns, not significant. Data are reported as mean ± SEM.

Four weeks after infecting GABA^CCK^ neurons with AAV (AAV9-mDLX-DIO-Chronos-mCherry and AAV9-mDLX-DIO-mCherry as control) in the AC of CCK-Cre mice (**Figure 5D-E**), the mice underwent behavioral training to associate sound and shock memory. As shown in **Figure 5F**, we observed that freezing proportion in CCK-Cre mice was significantly lower than that in controls after 1 hour for consolidation (**Figure 5F**, two way ANOVA, Bonferrori adjustment, F_2,12_ = 10.30, p = 0.002; Control: Pre 21.04 ± 4.18 % vs Post 87.21 ± 6.62 %, p < 0.001; CCK-Cre mice: Pre 18.43 ± 7.12 % vs Post 50.94 ± 5.75 %, p < 0.001; Post_Control_ vs Post_CCK-Cre_, p = 0.002). Additionally, we analyzed freezing activities in PV-Cre mice expressing the same AAV. Interestingly, no discernible difference was observed between PV-Cre mice and controls, suggesting that activation of GABA^CCK^ neurons, rather than PV interneurons, attenuates sound-shock associative memory (**Figure 5E**, PV-Cre mice: Pre 17.26 ± 3.72 % vs Post 69.92 ± 4.46 %, p < 0.001; Post_Control_ vs Post_PV-Cre_, p = 0.16; Post_CCK-Cre_ vs Post_PV-Cre_, p = 0.03). This result indicate that selective activation of GABA^CCK^ neurons significantly impaired the animal’s sound-shock associative memory at the behavioral level.

## Discussion

Inhibitory neurons play a crucial role in regulating neuronal excitability and long-term synaptic plasticity, subsequently controlling the activities of neural circuits and influencing an animal’s physiological states from multiple perspectives^31,32^. One important function of inhibitory neurons is to modulate the firing rate of their excitatory counterparts^33^. Previous studies have demonstrated that repetitive firing or high-frequency stimulation of inhibitory neurons (GABAergic neurons) elicits neuroplasticity of inhibition, consequently inducing long-term potentiation of inhibitory synapses (i-LTP) in post-synaptic neurons in several key brain regions, including the neocortex and hippocampus^34,35^. Interestingly, our result also elucidated that activation of GABAergic neurons augments the inhibitory effects on pyramidal neurons in AC area via optogenetic manner. Additionally, we further unveiled that HFLS of the mDLX-Chronos infected neurons enhances the amplitude of spontaneous IPSC, presumably due to the modification in post-synaptic activities. For instance, newly expressed GABA_A_ receptors in the post-synaptic membrane could prolong inhibitory neuroplasticity in the hippocampus, as shown in earlier studies^36^. Concurrently, other forms of cortical interneuron-mediated i-LTP may be involved in the induction of i-LTP, including NMDA receptors^37^, brain-derived neurotrophic factor (BDNF) receptors^38^, as well as GABA_B_ receptors^39^. Our recent work uncovered that a novel CCK receptor (GPCR-173), rather than CCK-1 or CCK-2 receptor mediates i-LTP formation in the AC area^21^. Nevertheless, more pharmacological investigations are needed to elucidate the neuronal mechanisms underlying the i-LTP in future studies. such as identifying the synaptic receptors and signaling pathways that modulate inhibitory potentiation of GABA neurons in the AC region.

Recent studies have demonstrated that excitatory CCK projections from the entorhinal cortex (EC) facilitate the formation of heterosynatic LTP in the AC area ^19,22^. Coincidentally, inhibitory CCK is broadly expressed in the AC, although its neurobiological functions in modulating synaptic plasticity are still poorly understood. In this study, we uncovered that HFLS of Chronos expressing GABA^CCK^ neurons is necessary for generating i-LTP in pyramidal neurons in the AC area, and post-synaptic modifications are also required for this inhibitory potentiation. Besides, in the hippocampus, pre-synaptic synapses from PV interneurons or SST interneurons can undergo unique forms of neuroplasticity mediated by activation of the T-type voltage gated Ca^2+^ channel (VGCC) ^40^. These forms of i-LTP suggest that different neuronal mechanisms are involved depending on brain regions, neuronal connections and brain activities. Based on these investigations, we further examined the effects of these two interneurons on shaping the plasticity in the AC under the same manipulations. However, HFLS was unable to potentiate the IPSC both in the PV-expressing synapse and SST-expressing synapse. This finding suggests that GABA^CCK^ neurons exhibit different neuronal activities in regulating the post-synaptic firing compared to PV interneuron and SST interneurons in the AC area. Additionally, to precisely conclude the neuronal influence of inhibitory system on pyramidal neuron in the AC area, more protocols are needed to examine the inhibitory effects of PV interneurons or SST interneurons on target excitatory neurons, such as utilizing different frequencies of light pulses, different frequency of light burst and pairing patterns.

According to previous findings that endogenous CCK can be released via external high-frequency stimulations from vesicles in the presynaptic synapses, effectively strengthening pre- and post-synaptic connections^17^. Hence, we continuously explored whether neuropeptide CCK could facilitates the i-LTP in PV-expressing synapses and SST-expressing synapses of PV-Cre mice and SST-Cre mice, respectively. Intriguingly, HFLS of the Chronos-expressed synapses paired with exogenous CCK-8s notably increased the amplitude of IPSC in PV-Cre mice. Moreover, this inhibitory potentiation was also observed in the LFLS group (**Figure 6**). These results indicate that selective activation of PV-expressing projections by light stimulation with neuropeptide CCK are two essential conditions for inducing the i-LTP in the AC area. Furthermore, we also utilized the same protocol to SST-expressing synapses in SST-Cre mice and observed similar results in both HFLS and LFLS groups with the addition of CCK-8s during IPSC recording. Collectively, we deduce that CCK plays a crucial role in shaping the neuroplasticity of the inhibitory system in the AC area.

**Figure 6.**
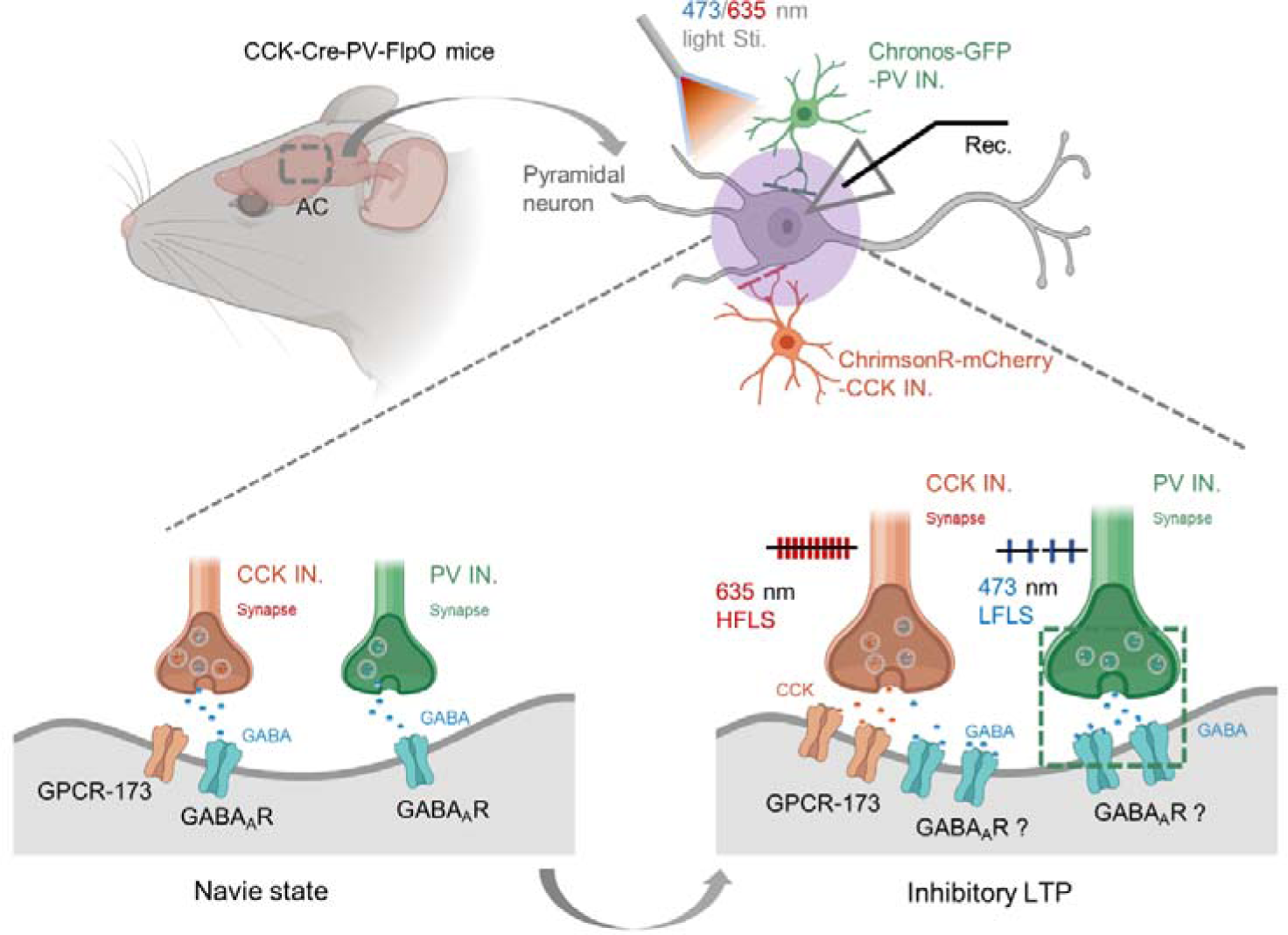
Schematic illustration depicts a possible cellular mechanism that HFLS of GABA^CCK^ release via by both postsynaptic GPCR173 receptor and GABA_A_ receptors produces LTP and impairs sound-shock associative memory. HFLS of Chronos expressed GABA^CCK^ neuron pairs with LFLS of ChrimsonR infected PV neuron triggers CCK released from the presynaptic terminals that facilitates the potentiates the inhibitory effects of both interneurons on post-synapses of pyramidal neurons in the AC area.

To mimic the effects of the endogenous CCK on other type interneurons in modulating neuroplasticity, we developed CCK-Cre-PV-FlpO mice to precisely manipulate the GABA^CCK^ neurons and PV interneurons simultaneously via dual-opsin (ChrimsonR and Chronos) stimulation techniques. Intriguingly, HFLS of Chronos-infected GABA^CCK^ neuron significantly enhanced IPSC in both GABA^CCK^ and PV synapses, while HFLS of ChrimsonR-expressed PV interneuron failed to elicit i-LTP formation in in either synapse. Since neuroplasticity activity has been considered as one of the major cellular mechanism for regulating the animal’s physiological states^41^. Therefore, we implemented a behavioral paradigm to test the influence of GABA^CCK^ neuron activity on sound-shock associative memory. We noticed that HFLS of GABA^CCK^ neurons effectively attenuated sound-shock associative memory compared to control mice and PV mice. Additionally, Whissell et al. found that activation of GABA^CCK^ neurons rich in cannabinoid type 1 receptor (CB1Rs) in the hippocampus ameliorates the cognitive impairment^42^. These results indicate that the neurofunctions of GABA^CCK^ interneurons vary depending on brain regions in behavioral aspects. Collectively, this study demonstrated that GABA^CCK^ neurons play an important role in neurofunctions at electrophysiological and behavioral levels, providing novel insights and future perspectives for understating the development of neurological diseases caused by disturbance in the E-I balance in the CNS.

### Materials and Methods Animals

All mice were bought from the Jackson Laboratory. For the in vitro brain slice recordings, immunohistochemistry and behavioral task, the following transgenic mice were utilized: CCK-ires-Cre (C57 background), PV-ires-Cre (C57 background), Sst-IRES-Cre (Sst^tm^^2.1(cre)Zjh^/J, C57BL/6 x 129S4/SvJae) Pvalb-2A-FlpO-D (C57 background). The animals were housed in a 12h-light/12h-dark cycle with food and water ad libitum at a stable temperature about 23-25°C. We adopted the standard procedures and the recommended primers for genotyping. All experimental methods and animals used in this study followed animal care, and biosafety guidelines were approved by the Animal Subjects Ethics Sub-Committees of the City University of Hong Kong.

### Virus injection Viral constructs

AAV9-mDlx-DIO-Chronos-mCherry-WPRE-pA and AAV-EF1α-fDIO-ChrimsonR-EYFP-pA were constructed and produced in the BrainVTA (China).

### Stereotaxic viral injection surgeries

Eight-week-old mice were used in this study. The mice were deeply anesthetized with pentobarbital (100 mg/kg) from Ceva Sante Animale Co., France, and supplemented with atropine (0.05 mg/kg) from Sigma, US. The depth of anesthesia was confirmed by the loss of muscular reflex to tail pinching. To maintain body temperature, a heating pad from RWD Life Science, China, was used to keep the mice at 35-37 °C.

Microinjections were performed using a microinjector 119 from World Precision Instruments, USA, and a glass pipette from World Precision Instruments, USA. The coordinates for targeting the auditory cortex (AC) were 4.5 mm anterior-posterior (AP), 2.5 mm medial-lateral (ML), and 3.5 mm and 7.0 mm dorsal-ventral (DV) relative to bregma. After the AAV injection, the mouse’s skin was sutured with sterilized sutures, and antibiotic ointment was applied to the incision to prevent bacterial infection and promote wound healing. The mice were then placed on a heated blanket until they regained consciousness. Finally, the mice were returned to the laboratory animal research unit for regular care.

### Brain slice preparation and patch-clamp recordings

Acute brain slices were prepared using a protective cutting and recovery method to achieve a higher success rate for patch-clamp^43^. Briefly, mice were anesthetized by intraperitoneal injection of pentobarbital sodium and transcardially perfused with 25-30 ml NMDG-ACSF (92 mM NMDG, 2.5 mM KCl, 1.25 mM NaH_2_PO4, 30 mM NaHCO_3_, 20 mM HEPES, 25 mM glucose, 2 mM thiourea, 5 mM Na-ascorbate, 3 mM Na-pyruvate, 0.5 mM CaCl_2_·4H_2_O and 10 mM MgSO_4_·7H_2_O; pH 7.3-7.4) saturated with 95% O_2_ and 5 % CO_2_ in prior. Slides containing the AC were transferred into NMDG-ACSF for 5-10 min at 32-34 °C to allow protective recovery and then transferred to room-temperature HEPES-ACSF (92 mM NaCl, 2.5 mM KCl, 1.25 mM NaH_2_PO_4_, 30 mM NaHCO_3_, 20 mM HEPES, 25 mM glucose, 2 mM thiourea, 5 mM Na-ascorbate, 3 mM Na-pyruvate, 2 mM CaCl_2_·4H_2_O and 2 mM MgSO_4_·7H_2_O; pH 7.3-7.4) for at least 1 h before recording.

After incubation, brain slices were transferred into the submerged recording chamber and continuously perfused (2-5 ml/min) with normal-ACSF (119 mM NaCl, 2.5 mM KCl, 1.25 mM NaH_2_PO_4_, 24 mM NaHCO_3_, 12.5 mM glucose, 2 mM CaCl_2_·4H_2_O and 2 mM MgSO_4_·7H_2_O) oxygenated with 95% O_2_ and 5% CO_2_. Slices were held in position by placing a fine-mesh anchor on the surface of the tissue. Whole-cell recordings were made at the AC and other corresponding brain regions using a Multi-clamp 700B amplifier and a Digital 1440A digitizer (both from Axon Instruments, Molecular Devices). The acquired signals were filtered at 2 kHz and digitized at 20 kHz. Slice quality was visualized on an upright fixed-stage microscope with a water immersion objective (Olympus BX51WI Fixed stage Upright Fluorescence Microscope IR-DIC). Patch-clamp pipettes with resistance between 3-6 mega ohms (MΩ) were pulled from borosilicate glass (WPI) using a Sutter-87 puller (Sutter). The pipettes were backfilled with internal solution (K-gluconate pipette solution), which contained 145 mM K-gluconate, 10 mM HEPES, 1 mM EGTA, 2 mM Mg-ATP, 0.3 mM Na**2**-GTP, and 2 mM MgCl_2_; pH 7.3; 290-300 mOsm. For the optogenetics manipulation, the laser generator (Inper, China) was connected to the whole-cell recordings system and the parameters of laser stimulation were controlled by the operation software of the laser machine.

### Behavioral task

Only male mice at 8 weeks of age were used in this behavioral task. AAV9-mDLX-DIO-ChrimonR-mCherry was injected into CCK-Cre mice and PV-Cre mice, while AAV9-mDLX-DIO-mCherry was used as a control.

On the first day of the experiment, all mice were placed in a square apparatus made of high-density polyethylene (25cm length, 25cm width, 25cm height) equipped with foot-shock and audio devices. The signals were automatically controlled by the TDT system. The mice were given one minute of free movement, followed by a tone cue and electrical stimulus (specific parameters are provided in Figure 5). After a 60-minute period for memory consolidation, 20 Hz light stimulation was delivered to the well-trained mice. Finally, on day 2, the mice were placed in a novel cylindrical apparatus (25cm length, 25cm width, 18cm height) and received a tone cue to assess their freezing response.

### Immunochemistry

For immunohistochemistry, the mice were deeply anesthetized with an overdose of pentobarbital sodium. They were then transcardially perfused with 30 mL of cold phosphate-buffered saline (PBS) followed by 30 mL of 4 % paraformaldehyde (PFA). The brain tissue was removed and post-fixed with 4 % PFA. It was then treated with 30 % sucrose in 4 % PFA at 4 °C for 2-3 days. The brain tissue was sectioned on a cryostat (60 μm for standard staining) and preserved with an antifreeze buffer at −20°C.

For immunostaining and analysis of co-labeled neurons in the auditory cortex (AC), serial sections containing the AC were selected. The brain sections were rinsed with PBS and blocked with a blocking buffer containing 10% goat serum in PBS with 0.2 % Triton X-100 for 2 hours at room temperature. The sections were then incubated with the primary antibodies (listed in Table 1) at 4 °C for 24 hours. After washing with PBS, the sections were incubated with the corresponding fluorophore-conjugated secondary antibodies (listed in Table 2) for 2.5 hours at 25 °C. The sections were then rinsed with PBS and stained with DAPI (1:5000 diluted in PBS) or mounted. For standard staining, the sections were mounted with 70 % glycerin in PBS on slides.

**Table 1.**
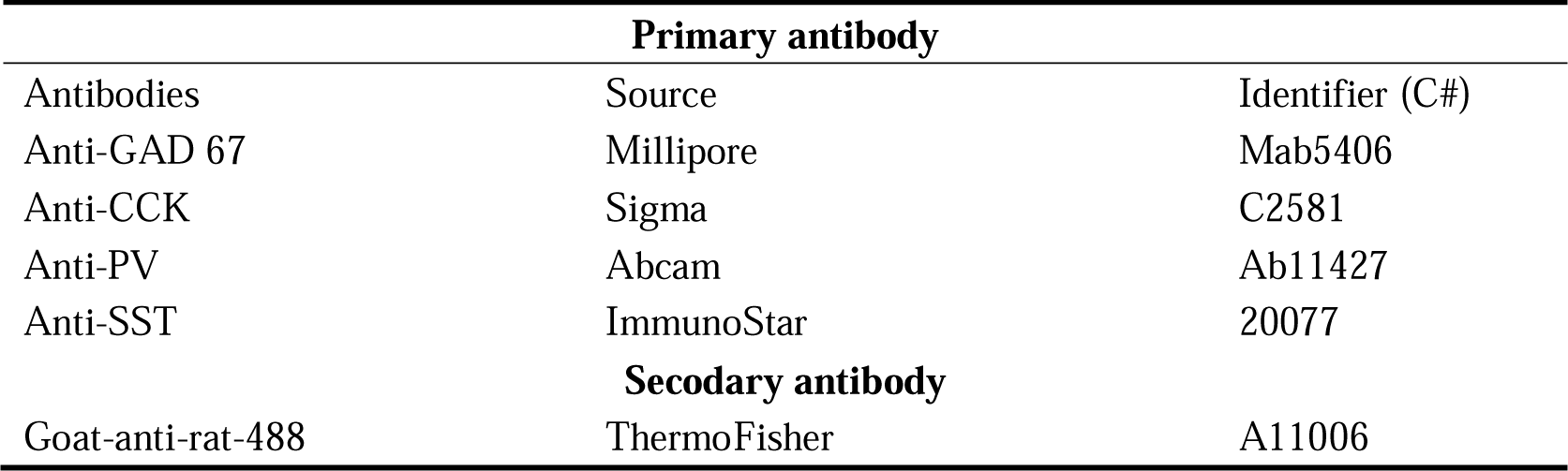
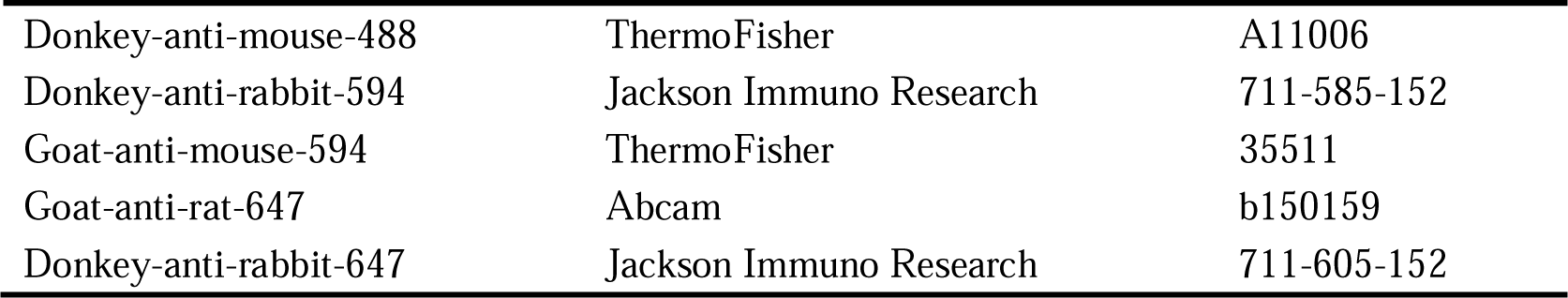
Primary antibody list for the immunostaining with brain tissues and cells.

Image acquisition was performed using an LSM 880 confocal microscope with 20X, 40X, 63X, and 100X magnification. The microscope was equipped with a time delay integration camera and performed line scanning to acquire the fluorescent signal at a high resolution.

### Data acquisition and analysis

All patch-clamp data were recorded using Axon Molecular Devices (Amplifier 700B and Digidata 1440A) and Clampex software. Except for the spontaneous events that were analyzed using the Mini Analysis Program, other patch-clamp data were processed using Clampfit 11. Group data are represented as mean ± SEM (standard error of the mean). SPSS 27.0 (IBM, Armonk, NY) was used for the statistical analysis (Two-way ANOVA, one-way ANOVA and pair-T test).

## Acknowledgments

This work was supported by funding from the following: Hong Kong Research Grants Council, General Research Fund: CityUHK 11101521, CityUHK 11103922, CityUHK 11104923, CityUHK 11104524. Hong Kong Research Grants Council, Collaborative Research Fund: C1043-21G. Hong Kong Research Grants Council, Theme-Based Research Scheme: T13-605/18-W. Hong Kong Research Grants Council, Senior Research Fellow Scheme: SRFS2324-1S02. Innovation and Technology Fund of the Hong Kong SAR, China: GHP_075_19GD. Hong Kong Health Bureau, Health and Medical Research Fund: 09203656, 08194106. Innovation Technology Commission of the Hong Kong SAR, China: Health@InnoHK program. We also thank the following charitable foundations for their generous support to J.H: Wong Chun Hong Endowed Chair Professorship, Charlie Lee Charitable Foundation, and Fong Shu Fook Tong Foundation.

## Authors contribution

JFH and GZ designed the study. GZ and KKP conducted the experiments. FWH, GZ KKP and XC analyzed the data. FWH, GZ and KKP created the figures. FWH and GZ wrote the manuscript. JFH edited the final manuscript.

## Ethics approval and consent to participate

Not applicable.

## Data availability

Data available on request from the authors.

---

**SFigure 1:**
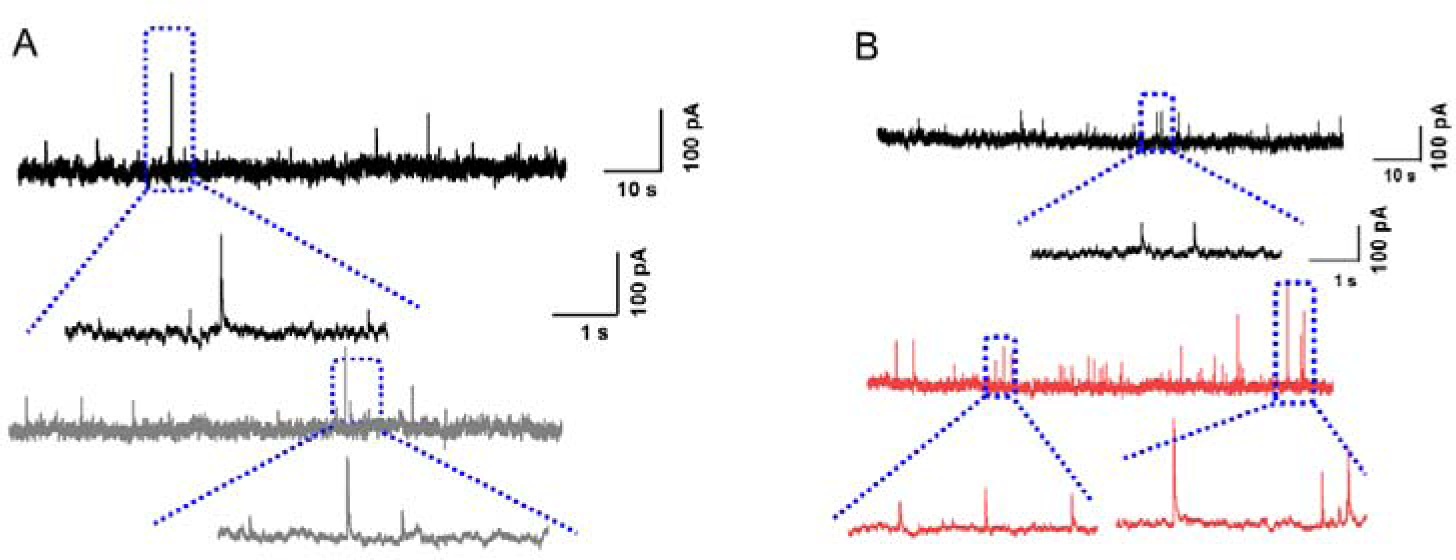
(A) Representative slPSC traces before (black) and after (grey) LFLS. (B) Representative slPSC traces before (black) and after (red) HFLS.

**SFigure 2.**
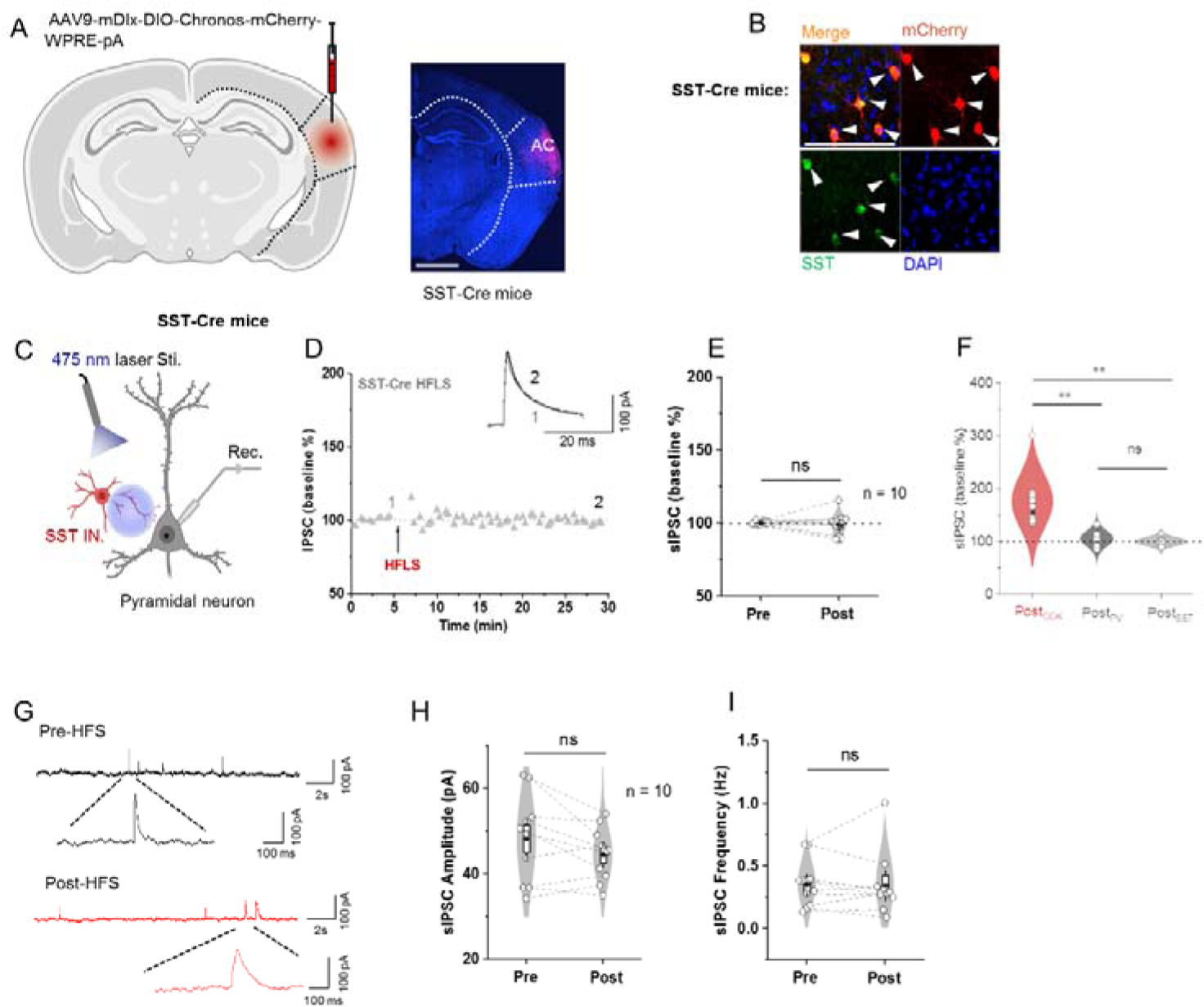
High frequency light stimulation ofGAJ3A^CCK^ neuron induced long-term potentiation in the inhibitory synapses of the GABA-Cre mice. (A) Schematic illustration of Cre-dependent AAV vector (D10-Chronos-mCherry) injection into the AC of SST-Cre mice to specifically infect GABAergic neurons, respectively. (B) Confocal image of virus expression in the AC of SST-re mice. White arrows indicate the colocalization of SST with mCheryy expressed SST positive neurons. Scale bar: 50 µ111. (C) Schematic of the IPSC recording of the SST interneuron in the A (D) Normalized amplitude of IP Cs before and after 635 nm wavelength HFL in S T-Cre mice. (E) Statistical comparison of the amplitL1des of IPSCs before and after 635 mn wavelength HFLS in ‘ST-Cre. (F) Statistical comparison of the amplitL1dcs of lPSCs before and after 635 nm wavelength HFLS in CCK-Cre, PV-Cre and SST-Cre mice. (G) Representative sIPSC traces of SST interneurons before (upper) and after (bottom) I!FLS in the SST-Cre mice. (H) Quantitative analyses of sIPSC amplitude before and after the *635* nm wavelength HFLS in SST mice (red). (I) Quantitative analyses of sIPSC frequency before and after 1he 635 11m wavelength HFLS in SST mice (red).

